# Motivational trade-offs in bumblebees

**DOI:** 10.1101/2022.02.04.479111

**Authors:** Matilda Gibbons, Elisabetta Versace, Andrew Crump, Bartosz Baran, Lars Chittka

## Abstract

Mammals can supress their nociceptive responses to prioritise other important responses via endogenous modulation from the brain. It is well established that insects display nociception, but not whether the insect brain can modulate nociceptive processing. To address this question, we investigated whether bumblebees’ (*Bombus terrestris*) attraction to higher sucrose solution concentrations reduces their avoidance of noxious heat. Bees were given the choice between either unheated or noxiously-heated (55°C) feeders with different sucrose concentrations. The feeders were associated with colour stimuli to act as conditioned cues. Bees fed more from higher sucrose concentration heated feeders than lower sucrose concentration unheated feeders. Further, bees’ “testing out” of feeders (landing but not feeding) reduced as the experiment progressed, demonstrating that conditioned colour cues informed the bees’ behaviour. Therefore, bees trade off competing conditioned motivational stimuli to modulate nocifensive behaviour, suggesting a form of pain perception.

## Introduction

Nociception and nocifensive behaviour – the detection of and response to noxious stimuli – occurs across many animal taxa, including insects (e.g. (1)). In mammals, neurons descending from the brain can facilitate or reduce nociception and nocifensive behaviour (2). The adaptive function of reducing nociception is to ensure that the subjective feeling of pain does not compromise the animal’s performance in acquiring another motivational requirement (2). For example, if an animal is food-deprived and sustaining injuries from fighting with its prey, reducing nociception and pain could improve fighting performance, and thus the chance of alleviating starvation. Fruit flies (*Drosophila melanogaster*) display such starvation-induced reduction in nocifensive behaviour (3). These findings illustrate that survival-linked states and stimuli can reduce nociception and nocifensive behaviour in both mammals and insects. For mammals, however, mental representations of states or stimuli can also drive this reduction. For example, imagining or expecting a state or stimulus that would reduce pain reduces nocifensive behaviour and pain in humans (4–6). It is unknown whether insects are also capable of this centrally-controlled reduction of nocifensive behaviour (7–9).

We tested whether a different motivational stimulus can modulate nocifensive behaviour in bumblebees (*Bombus terrestris*). We used a motivational trade-off paradigm, where animals must flexibly trade-off two competing motivations. For example, hermit crabs require higher voltages of electric shock to evacuate preferred *Littorina* shells than non-preferred *Gibbula* shells (10, 11). Shock avoidance is traded off against shell preference. We built on this paradigm by ensuring that the motivational trade-off relied on conditioned cues associated with the motivational stimuli.

## Results

Bees were given the choice of between high-quality feeders (containing 40% sucrose solution), and alternative feeders. In these alternative feeders, different groups of bees were offered 10, 20, 30 or 40% sucrose solution (Figure 3). Each of these groups experienced the unheated condition, with all feeders at room temperature (unheated), followed by the heated temperature condition, with the high-quality feeders at 55°C (heated) and the alternative feeders unheated. We also ensured that the sucrose concentration could not be detected unless the bee was feeding on the solution (sucrose solution has no scent and thus its concentration cannot be detected remotely (12)). We hypothesised that bees would avoid noxiously-heated feeders less when these dispensed higher sucrose concentrations relative to the unheated feeders.

When feeder types offered equal reward quality, bees avoided noxiously-heated feeders (mean proportion of feeding events on heatable feeder: 0.5 unheated, 0.3 heated; *z* = −4.050, *p* < 0.001, *N* = 10). Bees also preferred unheated high-quality feeders (40% sucrose solution) when alternative feeders contained lower sucrose concentrations (10%, 20%, or 30%; z = −13.12, *p* < 0.001, N= 41). However, the proportion of feeding events on 40% feeders reduced as the sucrose concentration of alternative feeders increased, and this effect was greater when the 40% feeders were heated (*z* = −2.068, *p* = 0.039, *N* = 32) (Figure 1). Further, the average number of landing events (“testing out”) on all feeders was significantly lower in the final foraging bout compared to the first (mean number of landing events on all feeders: 2.45 in first bout, 1.09 in final bout; *z* = −5.62, *p* < 0.001, *N* = 32).

**Figure 1.**
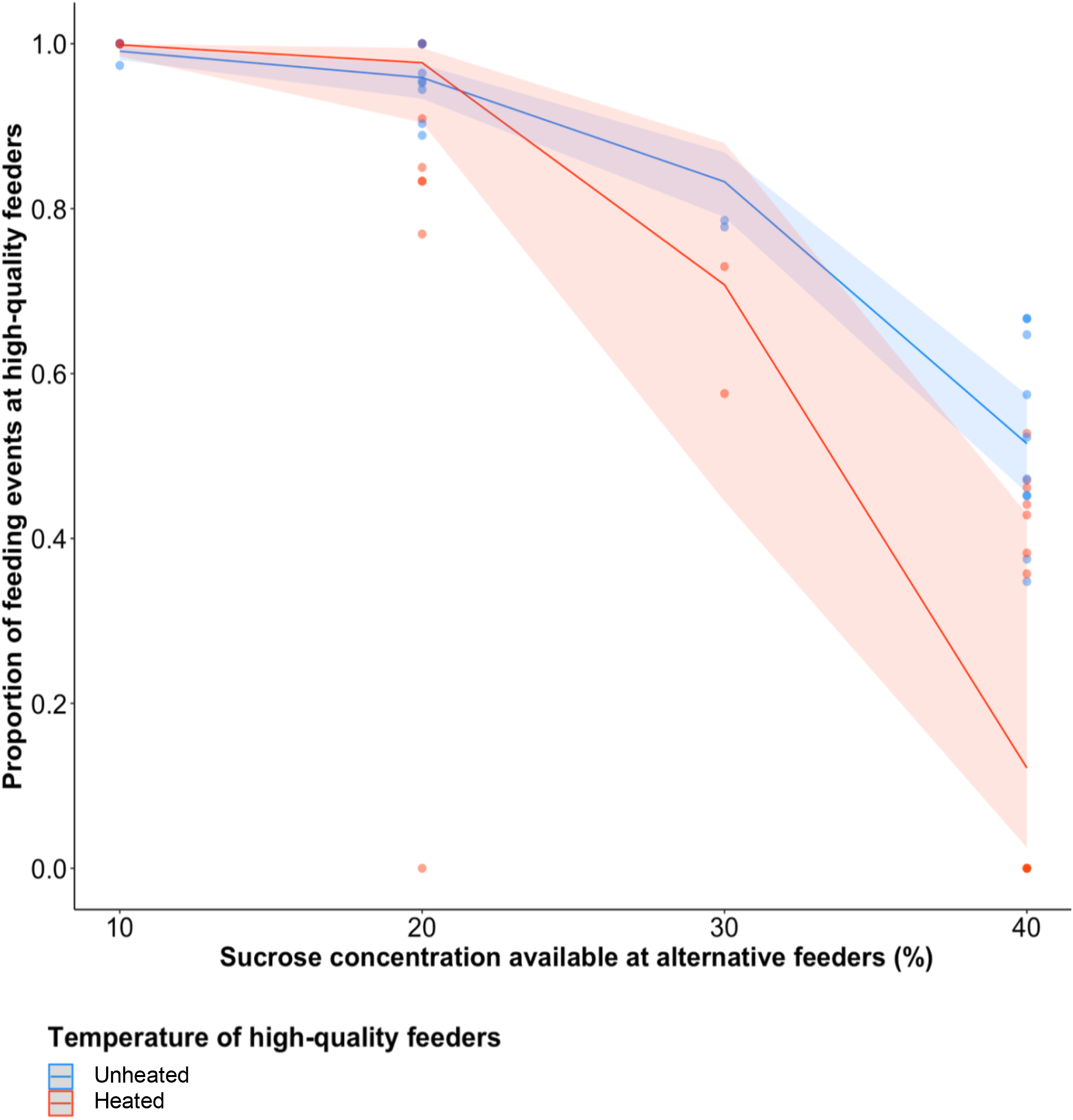
Proportion of feeding events on high-quality feeders in both temperature conditions and all concentration conditions. Shaded areas represent 95% confidence intervals (NS = non-significant; * = p < 0.05; ** = p < 0.01; *** = p < 0.001).

## Discussion

Bumblebees avoided heated feeders less when these dispensed higher sucrose concentrations than unheated feeders. Thus, bees traded off their motivation to avoid noxious heat against their preference for higher sucrose concentrations. The sucrose concentration could only be physically detected when the bee was directly on the unheated feeder, and since direct “testing out” of feeders decreased as the experiment progressed, we can conclude that the bees learned to associate the feeders’ contents with their colour cue and/or spatial location. This demonstrates that the trade-off relied on associative memories, rather than direct experience, of the stimuli.

Bees’ ability to trade off heat avoidance against sucrose preference indicates that conditioned motivational stimuli can influence nocifensive behaviour. This ability is consistent with the sensation of pain (8, 13–15), although, as in other animals, it is not conclusive proof. Further, attraction to conditioned cues for sucrose solution is processed in the insect brain (16, 17), so our results also suggest that the insect brain can modulate nociceptive behaviour endogenously, which was previously an open question for insects (9). Therefore, our results suggest that endogenous modulation and pain perception in insects are plausible and should be further explored.

## Materials and Methods

### Experimental set-up

Forty-one forager bees from eight colonies were tested. Bumblebee nests (Biobest, Belgium) were kept in wooden boxes connected to the testing arena, which contained four feeders (see Figure 2). Each feeder was on top of a heat-pad, and had four pink or yellow Perspex squares (25mm × 25mm) to act as conditioned stimuli (Figure 2). The feeders were arranged in a semicircle, 15cm apart and 30cm from the arena entrance, with alternating colours (Figure 2).

**Figure 2.**
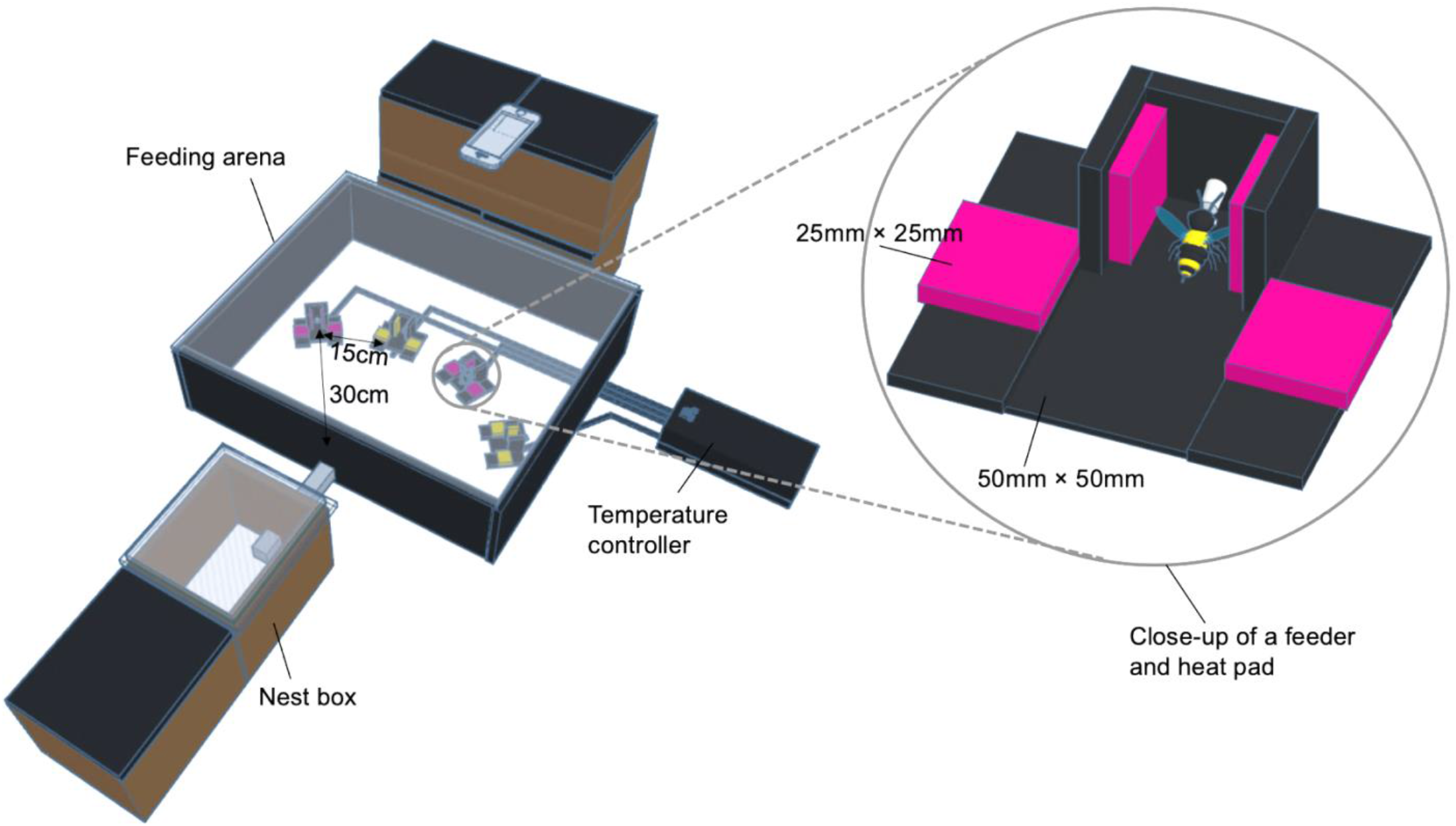
Testing arena set-up (not to scale).

**Figure 3.**
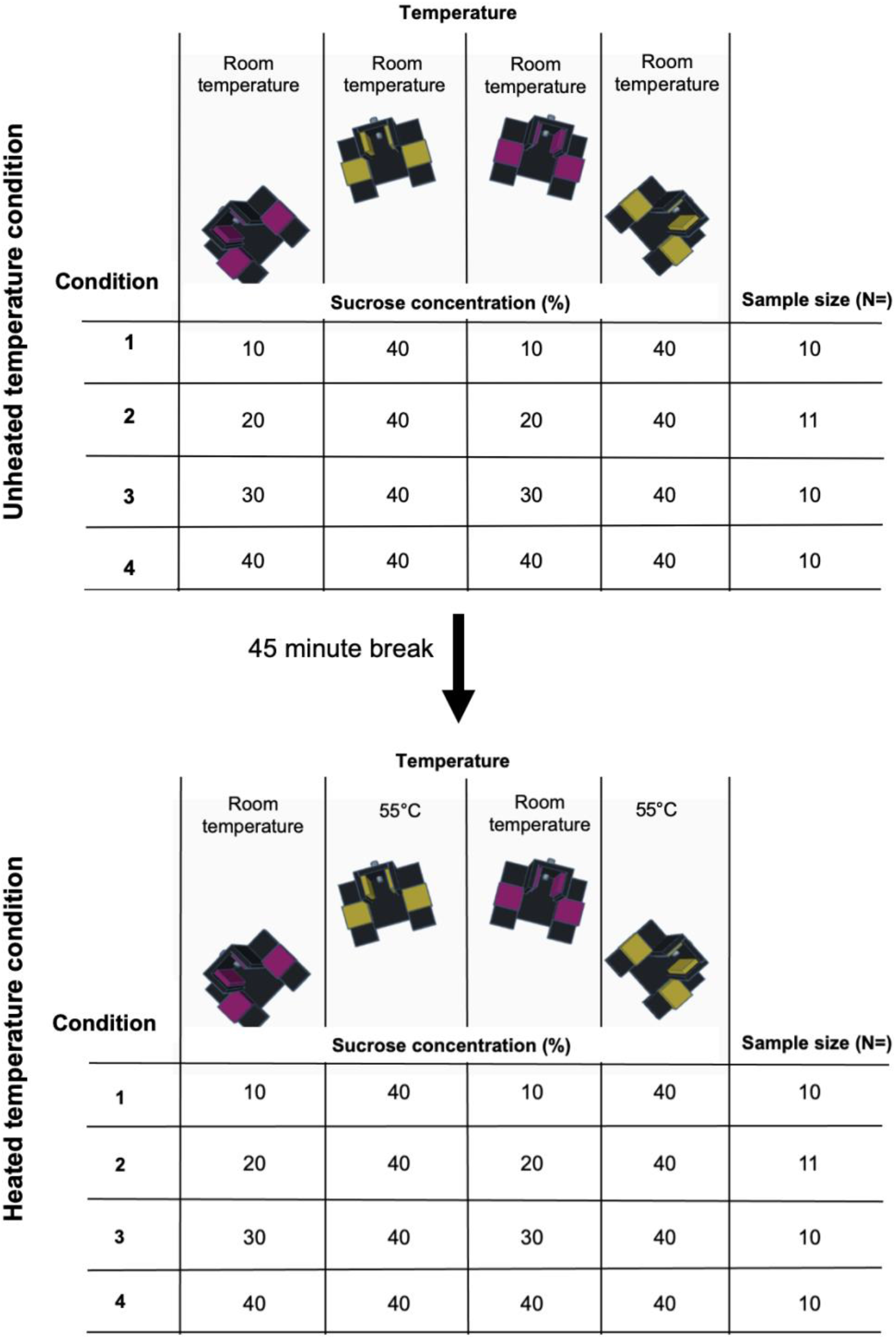
Flow chart depicting the order in which the temperature conditions were experienced, the temperature of the feeders in each temperature condition, and the sucrose concentration of the feeders in each concentration condition. Each of four different groups of bees (corresponding to one horizontal line in the tables) experienced only one combination of concentrations and temperatures in each phase. Color and positions of feeders were counterbalanced during the experiment.

### Training and testing

The bees first underwent training to familiarise them with the feeder conents (see supplementary information). Before testing, a forager bee was chosen, and only that bee was let into the arena. The bee was allowed to feed from the four feeders, which each contained 20μl of sucrose solution.

First bees underwent with an experimental condition where all feeders were unheated. Feeders were refilled every time the bee emptied them and landed on another feeder. Each bee completed 5-10 bouts (nest-arena-nest cycle) until it stopped foraging (no attempt to re-enter the arena for over 20 minutes). After a 45-minute break allowed the heat-pads to heat up, the bee entered the second testing stage (the heated temperature condition) where two of the four feeders were heated to 55°C (temperature was recorded using an infrared camera and infrared thermometer).

The same testing protocol was followed. Feeder colour and order were counterbalanced (high-quality feeders were pink for 20 subjects, and yellow for 21 subjects; the position of the feeders from left to right was ‘pink, yellow, pink, yellow’ for 18 subjects and the opposite for 23 subjects).

Throughout the heated and unheated conditions, the arena and feeders were cleaned with 70% ethanol every three bouts, as well as during the 45-minute break. We recorded the number of feeding events, defined as: “proboscis extended and reaching the sucrose-dispensing part of the feeder, and abdominal pumping visible for more than three seconds” and landing events, defines as: “enters cardboard chamber of the feeder and does not perform a feeding event” using an iPhone 6s (Apple, USA). For each bee, we then calculated the mean proportion of feeding events on the high-quality feeders across all her foraging bouts for each temperature condition, and the total number of landing events per first and final foraging bouts for each bee. We analysed the data in R (R Core Team, Cran-r-project, Austria, version 1.3.1093), using generalised linear mixed-effect models (see supplementary information). All study data and materials and methods are included in the article and supplementary information.

## Supplementary Information

### Animal care

Insects are not protected under animal welfare law in the United Kingdom, but we followed the Association for the Study of Animal Behaviour’s Guidelines for the Use of Animals in Research (1). Relatively few bees were tested (N = 41). Moreover, although the experiment involved noxiously-heated feeders, bees were never forced onto these – food was always available at unheated feeders.

Bees were kept in either 40×28×11 cm or 28×16×11 cm two-part wooden boxes connected via a Perspex tunnel (25 cm length; 3.5 × 3.5cm cross-section) to the testing arena (Perspex-roofed flight arena; 56cm × 56cm), which had four feeders (see Figure 2). One half of the two-part boxes that colonies were kept in contained the nest and had a wooden top to maintain 24-hour darkness. The other half had a 1cm-deep gravel substrate and a Perspex top, with a 12:12 hour light:dark cycle. A 1 cm-diameter hole connected the two sections. For ventilation, boxes also had four 2 cm-diameter exterior holes, which were covered with gauze to prevent escapes. Temperature was maintained at 23 °C. Colonies received seven grams of pollen (Natupol Pollen, Koppert Biological Systems, UK) every two days.

### Heat-pads

Heat-pads were custom-built using a single side copper-clad (copper thickness 70μm) standard printed circuit board laminate (made of woven fibreglass and epoxy resin, 1.57mm thick, 50mm × 50mm each) with an etched resistive heating element consisting of 18 loops (path width 0.6mm). The surface of each heat-pad was fitted with LM35 temperature sensors with thermally conductive adhesive (AG TermoGlue, TermoPasty, Poland). Heat-pads were powered with 12V DC, regulated by TIP122 transistors, driven with Arduino Uno Rev3. The temperature readouts were smoothed with a Kalman filter (SimpleKalmanFilter library).

### Training and testing

We identified foragers as bees that left the nest to feed more than four times in ~30 minutes, and marked these on the dorsal thorax with either an Opalith number tag (using Loctite Super Glue) or a coloured Uni POSCA marker pen. Before testing, we removed the bees from the arena using a plastic cup and a Perspex square to catch each bee and return her the nest. We then cleaned the arena and feeders with 70% ethanol solution. When tests were not being run, bees underwent group training, in which the whole colony could access cotton wool soaked with sucrose solution on the feeders in the arena. Then, after a forager was chosen, they underwent an individual training phase that consisted of the feeders not being refilled until the bee had consumed the sucrose solution from all four feeders, to ensure that she had experience of every feeder. During testing, if a forager spent five minutes in the testing arena without feeding, we returned her to the nest. Heat pad temperature was recorded using an infrared camera (FLIR One Infrared Camera, USA) and infrared thermometer (CASON CA380 Infrared Thermometer, TMS Europe)).

### Inclusion criteria

Our hypothesis relied on the bees distinguishing between the two concentrations of sucrose solutions and reliably choosing the higher one. The inclusion criterion for concentration conditions 10%, 20% or 30% was that the proportion of feeding events at the feeders containing 40% had to be significantly different from 0.5. The inclusion criterion for concentration condition 40% was that the proportion of feeding events at the feeders containing 40% had to not be significantly different from 0.5. One bee from the 20% condition and eight bees from the 30% condition did not meet our inclusion criterion and were excluded, giving a total sample size of 32 bees.

### Statistical analysis

We analysed the data in R (R Core Team, Cran-r-project, Vienna, Austria, version 1.3.1093), using generalised linear mixed effect models (GLMMs; packages: “lme4”, (2) “car” (3). We identified the most parsimonious models through the ANOVA function and stepwise backward elimination. As our data were proportions, and thus binary, we used a binomial distribution and logit link function. We checked model assumptions using histograms and Q-Q plots, and considered p < 0.05 significant.

Four separate GLMMs were fitted. In the first model, we tested whether bees preferred unheated feeders to heated feeders. The response variable was proportion of feeds on the heatable feeder; the fixed effect was temperature condition (heated, unheated); and the random effects were bee ID nested in colony ID. Model simplification removed location and colour of the heatable feeder and bout number. In the second model, we tested whether bees preferred 40% sucrose concentration feeders to lower-concentration alternative feeders. The response variable was the proportion of feeding events on the heatable feeder, and the fixed effect was concentration condition (10%, 20%, 30%, 40%). Model simplification removed location and colour of the heatable feeder, bout number, and colony ID. In the third model, we tested whether bees trade-off their heat aversion against their preference for higher sucrose concentrations. The response variable was proportion of feeding events on the heatable feeder; the fixed effects were temperature condition, concentration condition, and the temperature × concentration interaction; and the random effect was bee ID for the temperature condition only (because only the temperature condition was repeated measures). Model simplification removed location and colour of the heatable feeder, bout number, and colony ID. In the fourth model, we tested whether landing events increased significantly in the final bout from the first bout. The response variable was the number of landing events per bee; the fixed effect was bout; the random effect was bee ID.

